## Supplementary Information for "Dynamic allosteric networks drive adenosine A_1_ receptor activation and G-protein coupling"

### Table of contents

SI Figures……………………………………………………………….…………...3

SI Tables……………………………………………………………………………16

### SI Figures

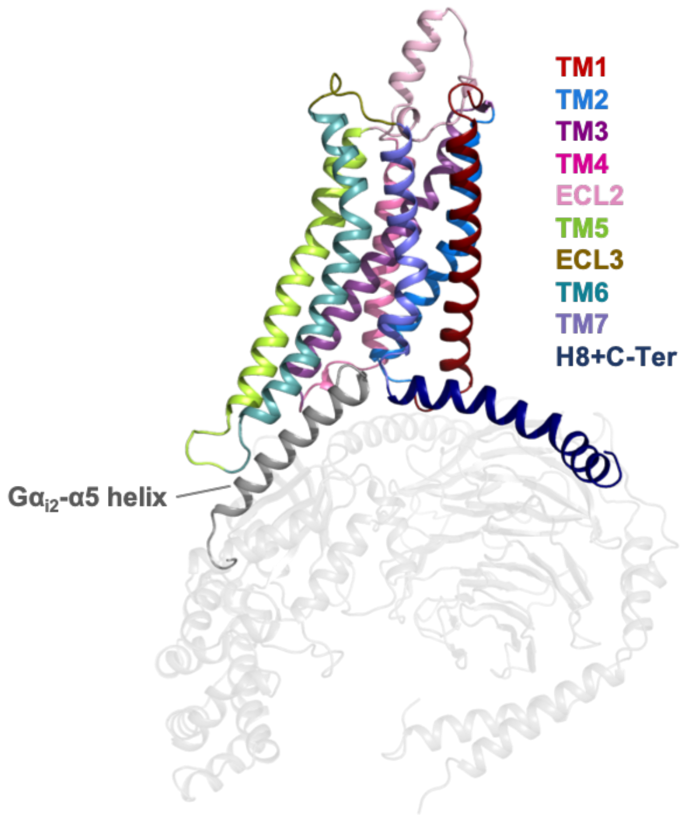


**Figure S1. Representation of the A$_1$R receptor in complex with heterotrimeric G$_{i2}$ protein (PDB 6D9H).** The helices and loops of the A$_1$R receptor are depicted with different colors while the heterotrimeric G$_{i2}$ protein in gray. The Gα$_{i2}$-α5 helix that binds into the intracellular cavity of the receptor upon activation is highlighted.

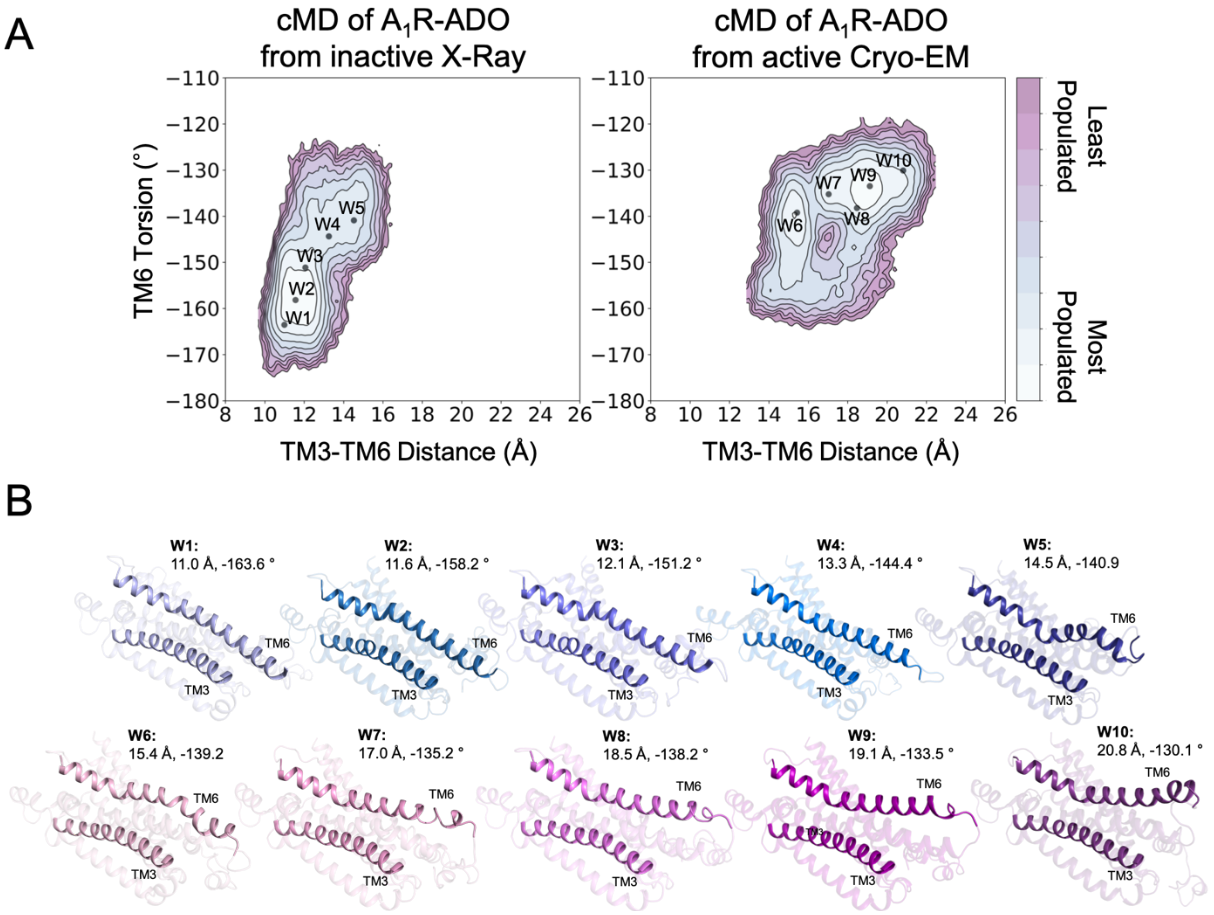


**Figure S2. Structures used as starting points for the walker metadynamics simulations. (A)** Population analysis of the $A_1R$ activation obtained from conventional molecular dynamics (cMD) simulations starting from the inactive X-Ray and active Cryo-EM structures, the coordinates of the walker structures (W1-10) are projected as black dots. **(B)** Representation of the walker (W1-10) structures. The CV1 (TM3-TM6 Distance) and CV2 (TM6 Torsion) values are shown. The TM3 and TM6 helices are highlighted.

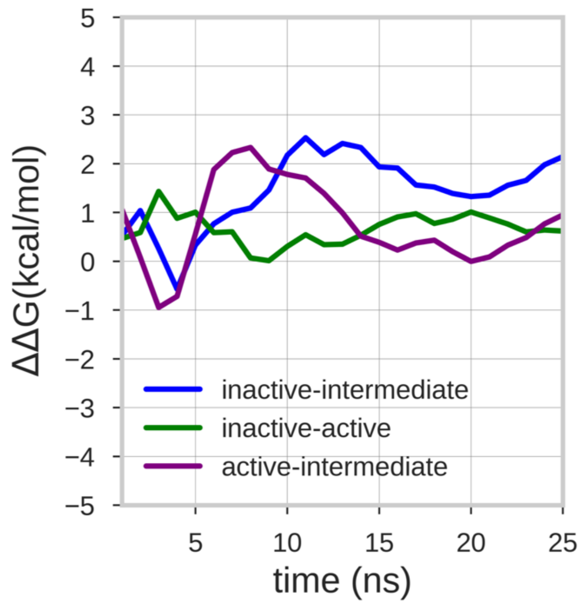


**Figure S3. Estimate of the free energy differences between the energy minima of the free energy surface.** The lines represent the mean ΔΔG value of the 10 walker replicas along the simulation time. The energy differences between the inactive-active, inactive-intermediate and active-intermediate energy minima are depicted in green, blue and purple, respectively.

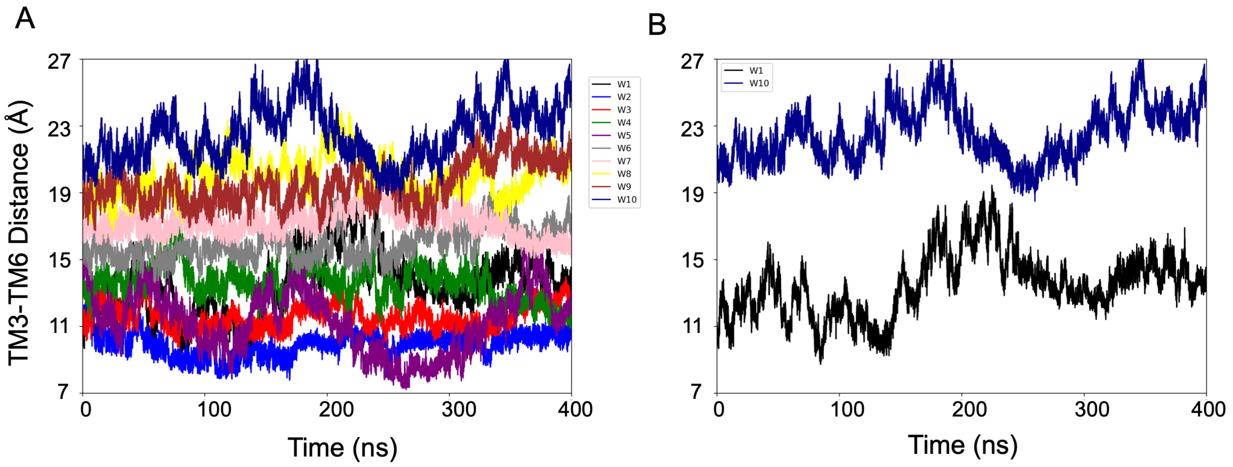


**Figure S4. Evolution of the CV1 (TM3-TM6 Distance) over the simulation time. (A)** Plot showing that the multiple-walkers (W1-10) sampling covers the CV space. **(B)** For clarity, only two walkers (W1 and W2) are represented.

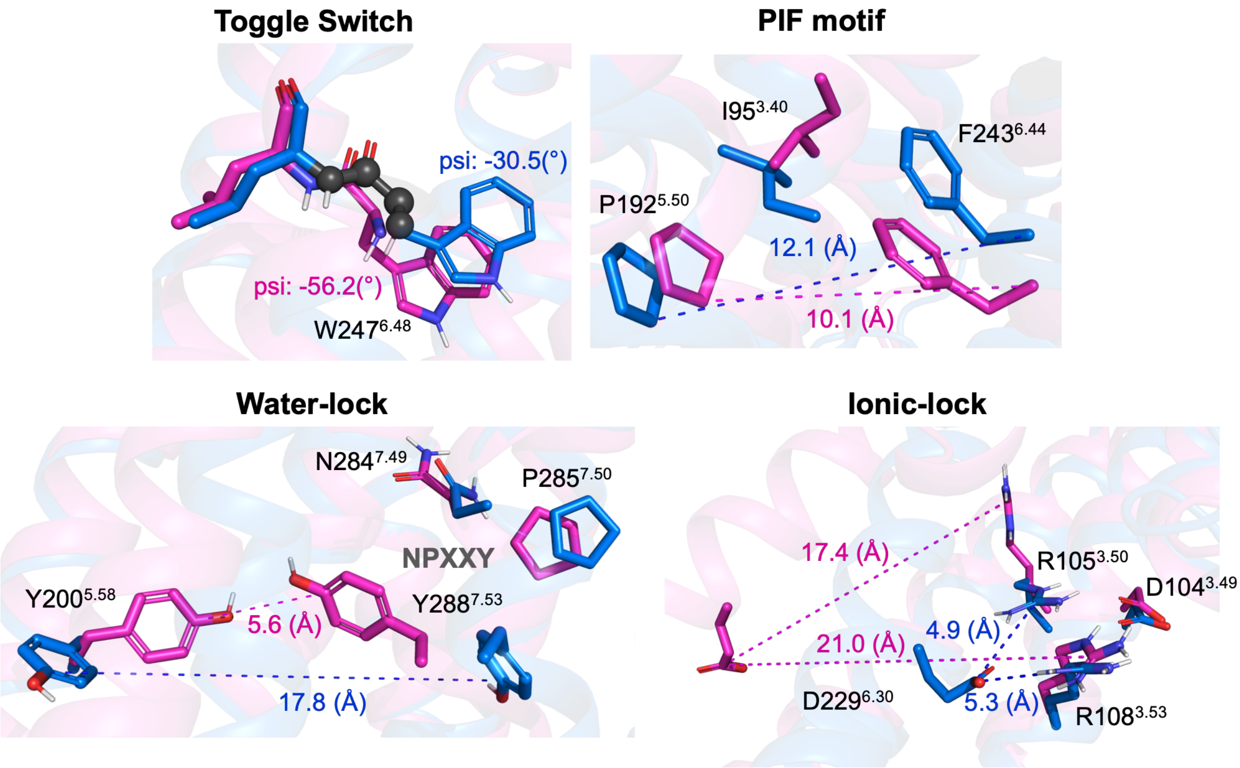


**Figure S5. Representation of relevant micro-switches for $A_1R$.** The inactive X-Ray (PDB 5N2S) and active Cryo-EM (PDB 6D9H) structures are displayed in blue and magenta, respectively. The metrics used in this work are highlighted in spheres for the psi dihedral and in dashed lines for the different distances. The values corresponding to both, inactive and active structures are also shown.

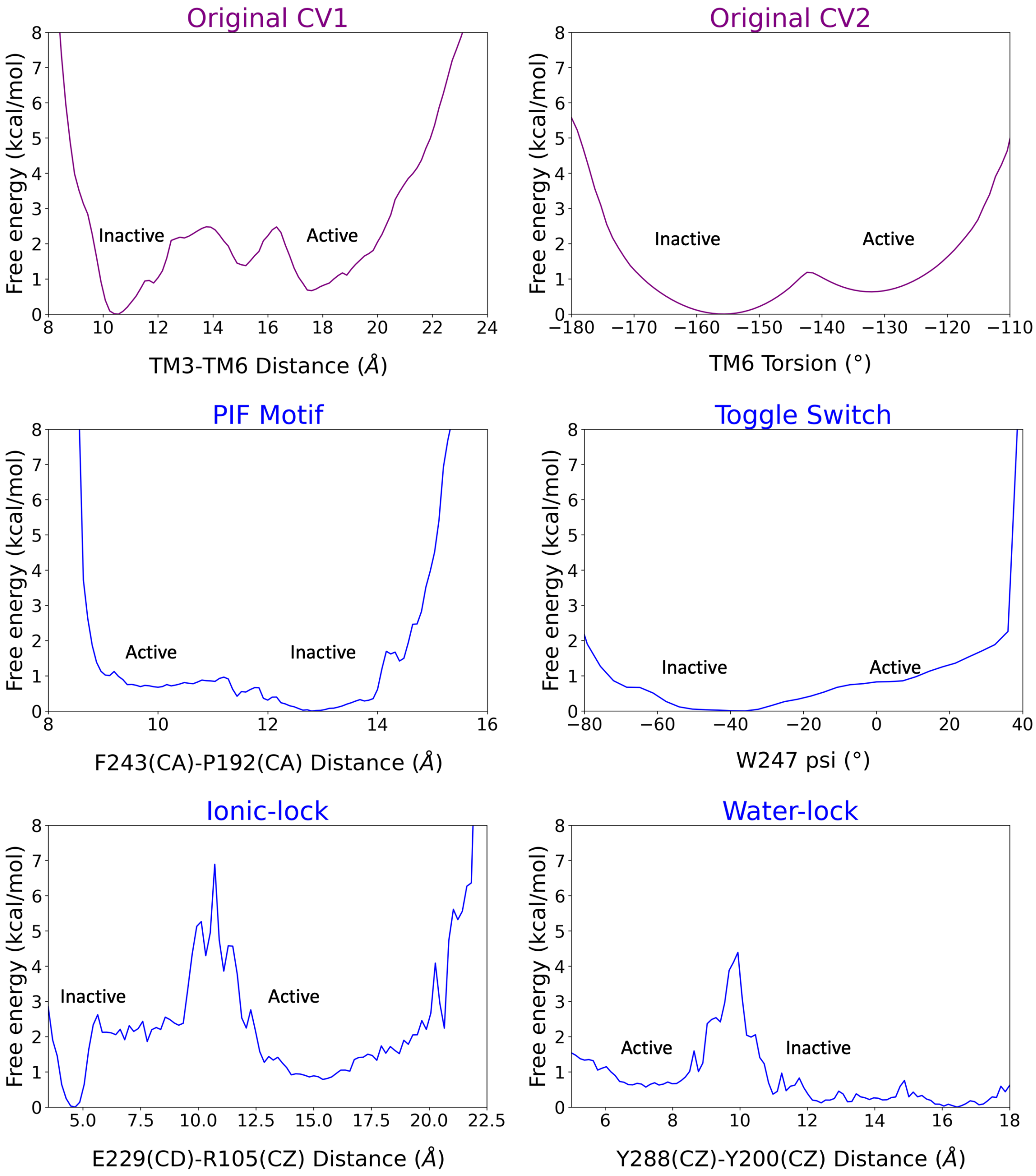


**Figure S6. Reweighting of the metadynamics simulations into 2D free energy profiles.** The original biased collective variables (CVs) are shown in purple while the unbiased CVs (i.e. the micro-switches) in blue. The inactive and active regions of the CV space are highlighted. Note that CV1 presents a higher contribution to the original energy barrier of activation than CV2. The PIF motif and toggle switch show similar energy barriers to CV2, while the ionic-lock and water lock display higher energy barriers than CV1. Capturing the distinct energy barriers associated with unbiased micro-switches highlights the accuracy of the metadynamics simulations to reproduce the pathway of activation.

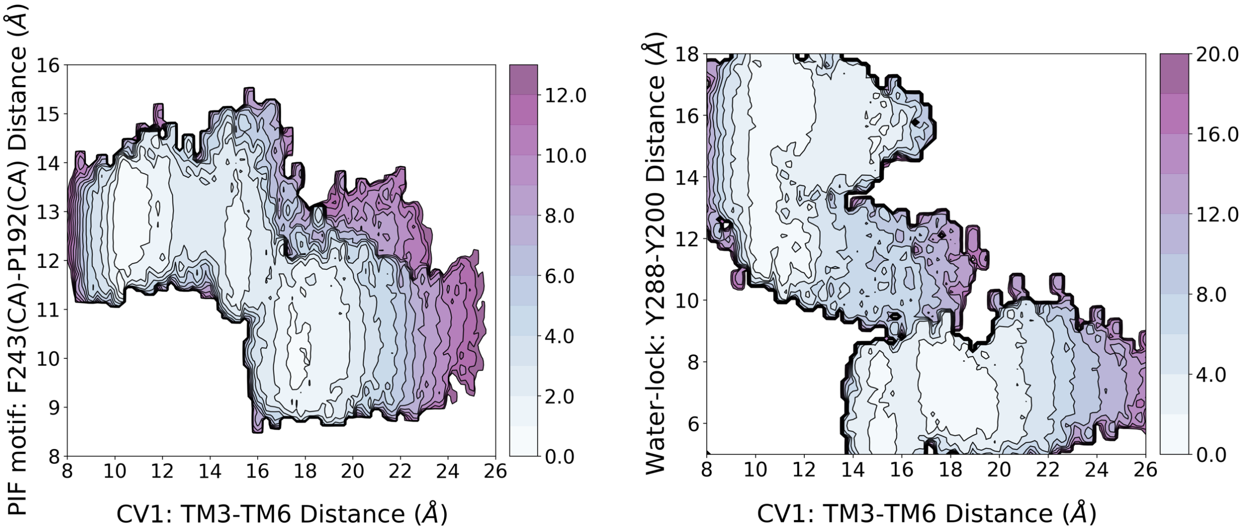


**Figure S7. Reweighting of the metadynamics simulations onto 3D free energy profiles.** The reweighting analysis shows that he activation energy barrier associated with the free energy landscape (FEL) of CV1 and the PIF motif is similar to that of the original biased FEL (i.e., CV1 and CV2). However, for the FEL associated with CV1 and the water-lock, the analysis predicts a higher energy barrier of approximately two-fold.

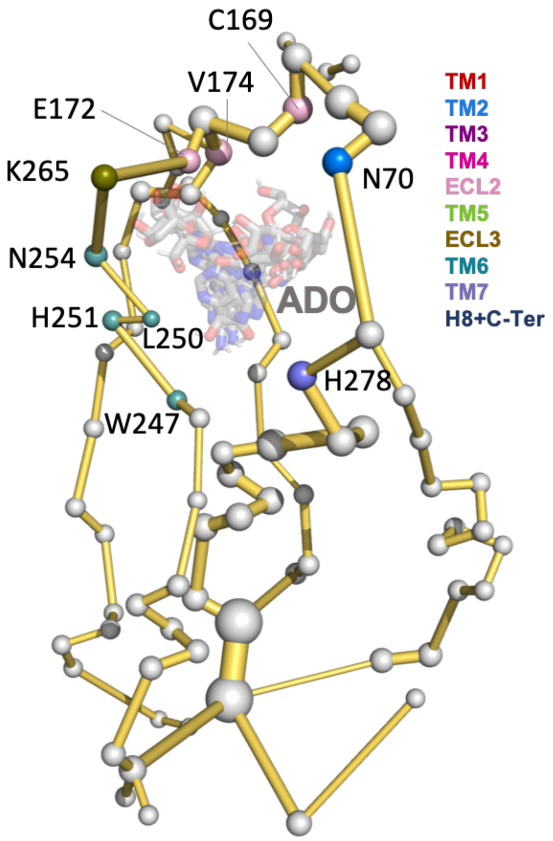


**Figure S8. Adenosine interactions with protein energy networks (PENs) residues of the $A_1$R-ADO conformational ensemble.** Representative structures from the broad range of conformations sampled by Adenosine (ADO) in the metadynamics simulations are shown as gray sticks. The PEN residues (nodes) that perform transient interactions with ADO are represented as colored spheres as a function of the receptor region (e.g. TM6 nodes in teal) while the rest of the nodes as white spheres. Note that the microswitch W247 is identified. The allosteric pathways (edges) are shown as yellow sticks.

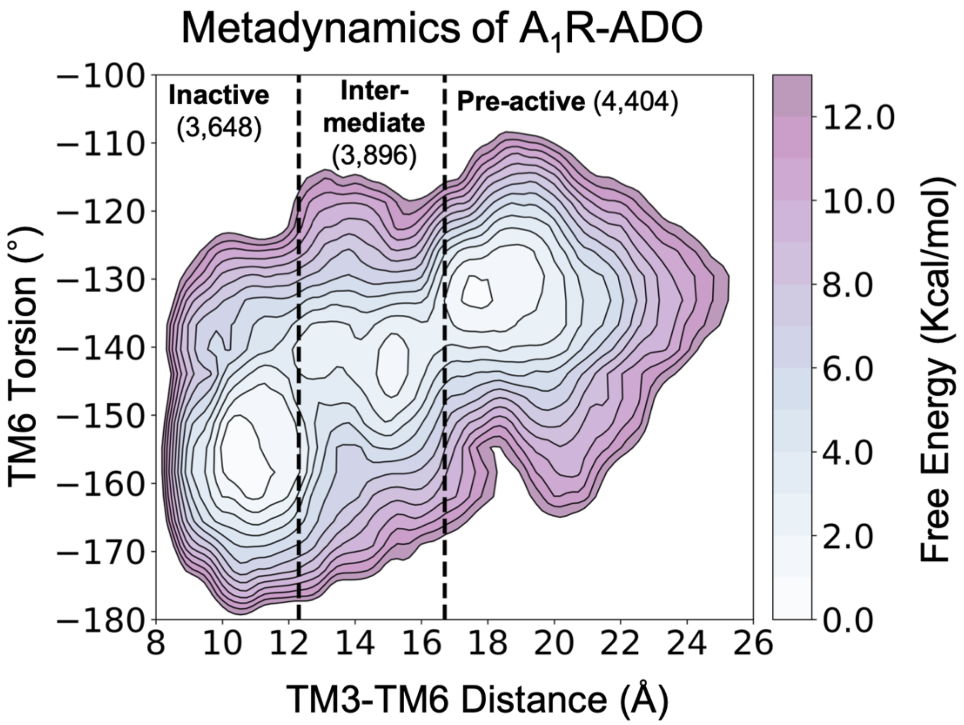


**Figure S9. Representation of the Free energy landscape (FEL) of $A_1$R-ADO activation split into conformational states.** The inactive, intermediate and pre-active regions of the FEL are separated by black dashed lines. The number of structures associated to each conformational state is also shown.

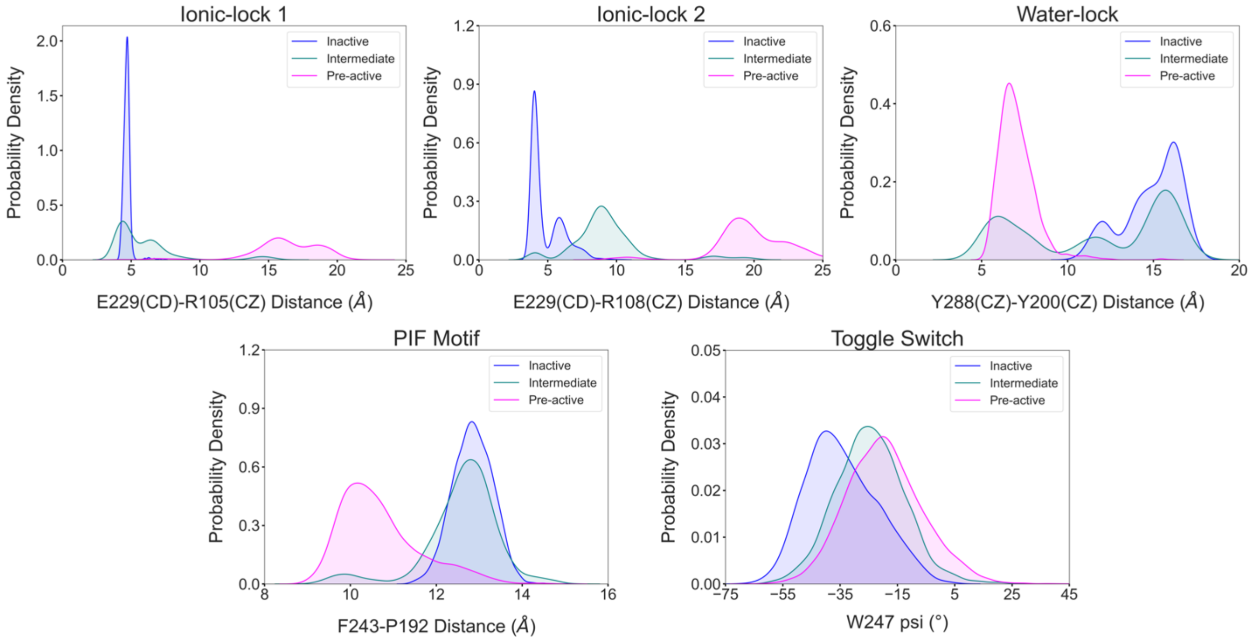


**Figure S10. Illustration of the conformational dynamics of the micro-switches along $A_1R$ activation.** The histograms corresponding to the inactive, intermediate and pre-active states are colored in blue, green and pink, respectively. The ionic-lock dynamics reveals that E229-R108 is the strongest interaction between TM6 and TM3 communication in the inactive ensemble, while in the intermediate state, the ionic lock is partially broken, and the strongest interaction shifts from E229-R108 to E229-R105. Notably, the micro-switches exhibit populations distributions that follow the progression of the receptor along the activation pathway in a correlated manner.

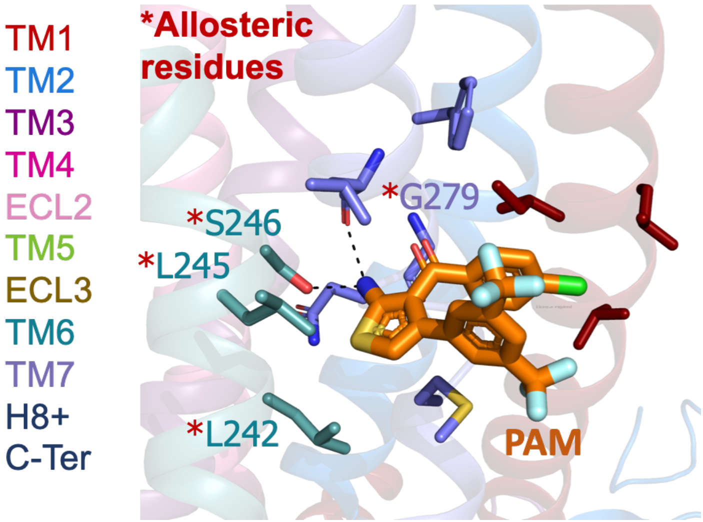


**Figure S11. $A_1R$ shallow pocket of the MIPS521 positive allosteric modulator PAM (PDB 7LD3).** The MIPS521 PAM is depicted in orange sticks while its interacting residues are colored as a function of the receptor region (e.g. TM6 nodes in teal). The experimentally identified allosteric residues are highlighted with a red asterisk.

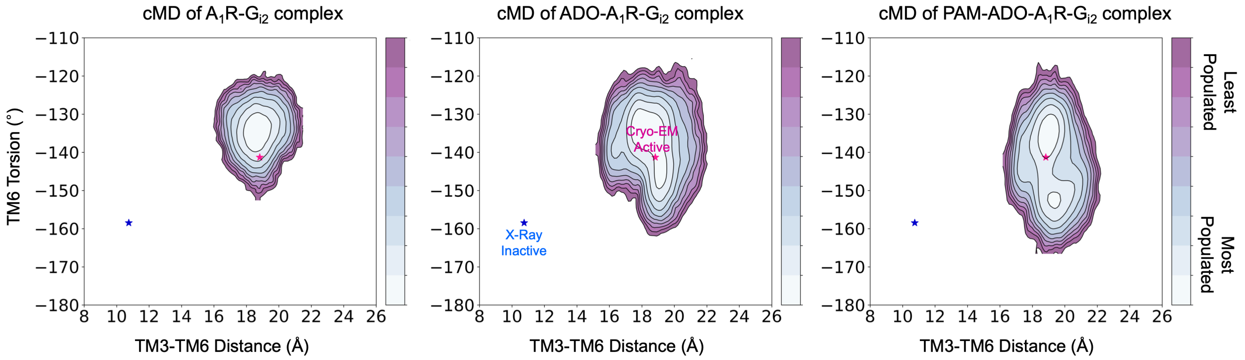


**Figure S12. Effect of ADO and PAM on the conformational landscape of $A_1R$ in presence of G-proteins.** Population analysis of $A_1R$ activation in the $A_1R$-$G_{i2}$ (left), ADO-$A_1R$-$G_{i2}$ (middle) and PAM- ADO-$A_1R$-$G_{i2}$ (right) complexes obtained from conventional molecular dynamics (cMD) simulations. The TM6 torsion corresponds to the dihedral angle formed by the alpha carbon atoms of L236$^{6.37}$, W247$^{6.48}$, T277$^{7.42}$ and T270$^{7.35}$. For the TM3-TM6 intracellular ends distance, we computed the center of mass (COM) distances between the backbone atoms of TM3(Y226$^{6.27}$, G227$^{6.28}$, K228$^{6.29}$, L230$^{6.30}$ and E230$^{6.31}$) and TM6(R105$^{3.50}$, Y106$^{3.51}$, L107$^{3.52}$, R108$^{3.53}$ and V109$^{3.54}$). The inactive and active X-ray and Cryo-EM coordinates are projected as blue and magenta stars, respectively. ADO and PAM induce additional flexibility to the TM6 torsion, which makes the Cryo-EM active coordinates more accessible.

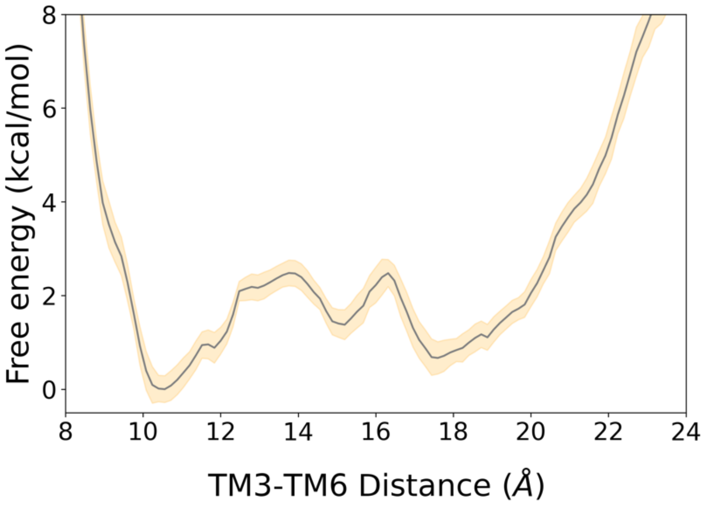


**Figure S13. 2D free energy landscape of $A_1R$ associated with the TM3-TM6 intracellular ends distance and its associated error.** The free energy landscape is represented by a gray line while its associated error corresponds to the yellow shaded area. The error was estimated using the block averaging technique, as described in the Materials and Methods section.

### SI Tables

#### Table S1. Description of the transient pockets location and containing Protein energy Network (PEN) residues in the inactive and intermediate states.

| Pocket | Location | Inactive (containing PEN residues) | Intermediate (containing PEN residues) |
|---|---|---|---|
| **PA** | ECL2 vestivule | **ECL2** (K173* and I167) | **ECL2** (K173*) |
| **PB** | Othosteric site + Secondary pocket | **ECL2** (E172, S176, I175, E170*, K168, C169, V174), **TM2** (G72, N70) **TM6** (N254), **ECL3** (K265*) | **ECL2** (E172, V174, I175), **TM7** (Y271, T277, I274, T270, H278) **ECL3** (K265*, P266) |
| **PC** | Extrahelical TM1-TM7 upper region | **TM1** (E16) | **TM1** (E16), **TM7** (Y271, I272) |
| **PD** | Extrahelical TM1-TM6-TM7 upper region | **TM1** (I19) | **TM1** (I19), **TM6** (L245*) |
| **PE** | Sodium ion site at the Internal channel | **TM2** (D55) | **None** |
| **PF** | Extrahelical TM2-TM3-TM4 lower region | **TM4** (A124, I128), **TM3** (D104), **TM2** (F47, S50) | **TM4** (A124), **TM3** (D104), **TM2** (F47, S50) |
| **PG** | Extrahelical TM3-TM5 lower region | **None** | **TM5** (E202), **TM3** (D104 and K110) |
| **PH** | Extrahelical TM5-TM6 lower region | **TM6** (A237, L236, L230, E229) | **TM6** (A244, F243,L240, A237, L236, L230, E229) |
| **PI** | Extrahelical TM6-TM7 lower region | **None** | **None** |
| **PJ** | Extrahelical TM1-TM7-H8 lower region | **TM1** (N27, L29, V30) | **TM1** (N27, L29, V30), **H8** (F229, R296) |
| **PK** | Extrahelical TM1-H8-C-ter lower region | **None** | **TM1** (W32 and A33), **H8** (K301, V297,T298, N293, K294) |
| **PL** | Intrahelical lower region | **TM3** (R108, R105), **TM6** (S235, K228), **TM2** (D42), **H8** (K294, D326) | **TM3** (R108, R105, I101, D104, V109), **H8** (N293, K294), **TM6** (S235, I236, I239), **TM2** (I48, D42) |
| **PM** | Extrahelical TM1-TM2-TM4 lower region | **TM2** (C46, V49, S50, D42), **TM4 (**A124, I128**), TM1 (**I31, V34**)** | **TM2** (S50, V49, D42, C46), **TM4** (A124, A125), **TM1** (I31, V34) |
| **PN** | Extrahelical TM5-TM6 lower region | **None** | **None** |

#### Table S2. Description of the transient pockets location and containing Protein energy Network (PEN) residues in the pre-active and fully-active states.

| Pocket | Location | Pre-Active (containing PEN residues) | Fully-Active (containing PEN residues) |
|---|---|---|---|
| **PA** | ECL2 vestivule | **ECL2** (K173*) | **ECL2** (K173*, I167 and W156*) |
| **PB** | Othosteric site + Secondary pocket | **ECL2** (E172, I175, E170*, K168, C169, V174), **TM2** (G72, I63), **TM7** (T27 **TM6** (W247, H251) **ECL3** (K265*, P266) | **ECL2** (E172, I175, E170*, K168, C169, V174), **TM2** (N70, G72), **TM7** (T270, Y271, A273, T277, H27 **TM6** (W247, L250, N254) **ECL3** (K265*, P266) |
| **PC** | Extrahelical TM1-TM7 upper region | **TM1** (E16, Y12, N9), **TM7** (Y271, I272) | **TM1** (E16, Y12, N9), **TM7** (Y271, I272) |
| **PD** | Extrahelical TM1-TM6-TM7 upper region | **TM6** (L242*, S246*), **TM7** (A282, M283) | **TM6** (L242*), **TM7** (G279*, M283) |
| **PE** | Sodium ion site at the Internal channel | **TM2** (D55), **TM6** (W247), **TM3** (S94), **TM7** (N284) | **TM2** (D55), **TM6** (W247), **TM3** (S94), **TM7** (N284, N280) |
| **PF** | Extrahelical TM2-TM3-TM4 lower region | **TM4** (A124, I128, C131), **TM3** (D104) | **TM3** (S93) |
| **PG** | Extrahelical TM3-TM5 lower region | **TM5** (E202, L195), **TM3** (D104 and K110) | **TM5** (E202), **TM3** (K110) |
| **PH** | Extrahelical TM5-TM6 lower region | **TM6** (A327, L236, E229) **TM3** (L201,Y200) | **TM6** (L240, A327, L236, I232, E229) |
| **PI** | Extrahelical TM6-TM7 lower region | **TM6** (L238, L242*) **TM7** (R291, M283, N284, I286) | **TM6** (L238, I239,L242*) **TM7** (R291, N284, M283, Y288) |
| **PJ** | Extrahelical TM1-TM7-H8 lower region | **TM7** (I286) | **TM7** (F290, I286), **H8** (R296) |
| **PK** | Extrahelical TM1-H8-C-ter lower region | **TM1** (N37, V36 and W32), **H8** (E317, D316, N293 and K294) | **None** |
| **PL** | Intrahelical lower region | **TM3** (R108, D104, V109), **H8** (N293, K294), **TM6** (I232), **TM7** (I292), **T** | **TM3** (R108, V109)**, H8** (N293)**, TM6** (I232)**, TM7** (I292)**, TM2** (D42)**, TM5** (R208) |
| **PM** | Extrahelical TM1-TM2-TM4 lower region | **TM2 (**D42**), TM4 (**A124, I128**)** | **TM2** (D42) |
| **PN** | Extrahelical TM5-TM6 lower region | **TM6** (E229, K228) | **TM6** (E229)**, TM5** (R208) |
